# Airway epithelial response to RSV is impaired in multiciliated and goblet cells in asthma

**DOI:** 10.1101/2023.03.16.532356

**Authors:** Aurore C. A. Gay, Martin Banchero, Orestes A. Carpaij, Tessa Kole, Leonie Apperloo, Djoke van Gosliga, Putri Ayu Fajar, Gerard H. Koppelman, Louis Bont, Rudi W. Hendriks, Maarten van den Berge, Martijn C. Nawijn

## Abstract

In patients with asthma, respiratory syncytial virus (RSV) infections can cause disease exacerbations by infecting the epithelial layer of the airways, inducing an innate and adaptive immune response. The type-I interferon antiviral response of epithelial cells upon RSV infection is found to be reduced in asthma in most -but not all-studies. Moreover, the molecular mechanisms that cause the differences in the asthmatic bronchial epithelium in response to viral infection are poorly understood.

Here, we investigated the transcriptional response to RSV infection of primary bronchial epithelial cells (pBECs) from asthma patients(n=8) and healthy donors(n=8). The pBECs obtained from bronchial brushes were differentiated in air-liquid interface conditions and infected with RSV. After three days, cells were processed for single-cell RNA sequencing.

A strong antiviral response to RSV was observed for all cell types present, from both asthma patients and healthy donors. Most differentially regulated genes following RSV infection were found in cells transitioning from basal to secretory. Goblet cells from asthma patients showed lower expression of genes involved in the interferon response. In multiciliated cells, an impairment of the signaling pathways involved in the response to RSV in asthma was observed, including no enrichment of the type-III interferon response.

Our results highlight that the response to RSV infection of the bronchial epithelium in asthma and healthy airways was largely similar. However, in asthma, the response of goblet and the multiciliated cells was impaired, highlighting the need for studying airway epithelial cells at high resolution in the context of asthma exacerbations.

**What is already know on this topic:** The airway epithelium response to RSV is altered in asthma. However, literature remains conflicted about the exact changes in the antiviral response, and the mechanisms causing these changes are yet to be found.

**What this study adds:** This study describes extensively the response of the bronchial epithelial cells (BECs) to RSV for both healthy subjects and asthma patients, at a single-cell resolution. It highlights the major overlap between healthy and asthma in the antiviral response to RSV. It allows the identification of specific genes and cell types that show a different behavior in response to RSV in asthma compared to healthy.

**How this study might affect research, practice or policy:** Our study indicates that goblet and multiciliated cells are the most relevant BECs to further investigate in the context of drug development for RSV-induced asthma exacerbation. It also suggests that focusing research on the cross-talk between the epithelial and the immune cells, or into investigating a potential delayed response in asthma would be the best way forward into understanding the mechanisms involved in the asthma response to RSV.

## Introduction

The prevalence of asthma, one of the major respiratory diseases worldwide, has increased in recent decades[1]. Asthma exacerbations remain a frequent cause for medical emergencies, and as such, a heavy burden on our healthcare systems[2]. In addition, frequent exacerbations result in subsequent asthma-associated loss of quality of life. Most asthma exacerbations are triggered by allergens or viral respiratory tract infection, with studies reporting respiratory syncytial virus (RSV) as an important viral cause for asthma exacerbations in adults[3].

The airway epithelium acts as the primary defense against pathogens, and therefore represents the primary target for RSV infection. In asthma, the airway epithelium is more susceptible for injury and displays a reduced barrier function. The airway epithelium in asthma has a decreased expression of E-cadherin and tight junction proteins, together with an increased basal cell proportion[4]. These changes of the airway epithelium are thought to contribute to the impaired barrier formation in asthma that was reported in *in vitro* cultured bronchial epithelial cells[5]. In addition, studies indicated that the receptors involved in viral sensing were altered for individuals with asthma[6]. In particular, a decreased expression of pattern recognition receptors (PRRs) was observed in both mild and severe asthma patients[7], contributing to an increased susceptibility to viral infection for people with the disease[8,9].

RSV binds to airway epithelial cells through the association of the viral attachment glycoprotein with glycosaminoglycans linked to transmembrane proteins at the cell surface. The viral entry is then facilitated by the viral fusion glycoprotein [10]. Though the main host cell attachment factor interacting with the viral fusion protein is thought to be the fractalkine receptor CX3C-chemokine receptor 1 (CX3CR1), several other proteins have been proposed to facilitate the RSV entry to the cell, such as, the intercellular adhesion molecule 1 (ICAM1), the epidermal growth factor receptor (EGFR) and nucleolin[10]. Subsequent release of the viral RNA into the cytoplasm will activate two main PRRs: the Toll-Like receptors and retinoic acid-inducible gene I like receptor family members[11]. This triggers an early innate immune response activating type I and type II interferon and the NF**-**kB pathway [9]. In air-liquid interface (ALI) cultures derived from primary bronchial epithelial cells (pBECs) from asthma patients, an exuberant inflammatory cytokine response was observed after RSV infection [4]. However, literature remains conflicted on the actual changes happening in the viral response to respiratory infection in the asthmatic airway. Several studies reported that type-I interferon production and the subsequent interferon response were reduced in the asthmatic airway after rhinovirus or RSV exposure, compared to airways from healthy controls[12-15]. However, others found that this antiviral response was preserved in pBECs in asthma[16], or delayed[17], compared to pBECs from healthy subjects. Those inconsistencies in the previous findings could be explained by differences in the response to RSV from the various cell types of the airway epithelium.

To this day, virally induced exacerbations of asthma remain difficult to prevent. Despite the importance of RSV in asthma exacerbations, the molecular mechanisms and cell type-specific responses that cause differences in the bronchial epithelial response to viral infection between healthy individuals and patients with asthma remain incompletely understood. Previous investigations of these mechanisms using bulk transcriptomics could not unveil cell specificity, and these unmeasured differences in cell type composition could partly explain the inconsistency observed in the current literature. In this study, we aimed therefore to compare the transcriptional response of pBECs in a cell-type specific fashion, between asthma patients and healthy donors to RSV infection. After establishing the characteristics of our culture model in healthy and asthma-derived samples at baseline, we investigated the transcriptional response to RSV in the healthy-derived cultures and compared it to that of the asthma-derived cultures.

## Methods

### Subjects

We included samples derived from bronchial brushes from patients with asthma and from healthy controls as previously described[18]. The healthy donors showed normal lung function, defined as forced expiratory volume in 1 s (FEV 1)/forced vital capacity (FVC) lower limit of normal (LLN), FEV1 >80% predicted, an absence of bronchial hyper-responsiveness (BHR) to methacholine (provocative concentration causing a 20% fall in FEV 1 (PC 20) methacholine mg·mL−1) and no allergies (see Table 1 for subject characteristics). The medical ethics committee of University Medical Center Groningen (UMCG) approved the study and all subjects gave written informed consent. Details about the patient cohort can be found in the supplementary material.

**Table 1:** Clinical characteristics of the study cohort.

|  | Asthma (n=8) | Healthy (n=8) | p-value |
| --- | --- | --- | --- |
| Age | 50.62 (9.56) | 56.12 (7.68) | 0.225 |
| Gender = male | 4 | 4 | 1.000 |
| Smoking = past smoker | 0 | 4 | 0.083 |
| FEV1 %predicted | 94.25 (14.55) | 114.12 (5.46) | 0.003 |
| FEV1/FVC % | 72.21 (8.53) | 79.29 (3.74) | 0.050 |
| PC20 Methacholine mg/mL | 2.19 (1.95) | 999.00 (0.00) | <0.001 |
| Blood eosinophil count *10E<sup>9</sup> /L | 0.17 [0.13, 0.27] | 0.15 [0.11, 0.19] | 0.429 |

### Culture of primary bronchial epithelial cells

Primary BECs were obtained from the bronchial brushes and differentiated in ALI cultures. During the fourth week of culture, cells were infected with RSV and were processed for scRNAseq 72hpi. Enzyme-linked immunosorbent assay (ELISA) was performed on the apical washes of the cultures as described in the supplementary material.

### Single-cell RNA-sequencing

#### Library preparation and sequencing

Each sample was incubated individually with 0.025μg TotalSeq-B hashtag antibodies (BioLegend, cat#: 394631, 394633, 394635 and 394637), according to the manufacturer’s recommendation, and 4000 cells from each sample (RSV or control) were combined in pools of 4 samples. Resulting cell suspensions were loaded and libraries were prepared according to standard protocol of the Chromium single-cell 3′ kit v3.1 with Feature barcoding antibodies (10XGenomics). Quality and concentrations of the different libraries were assessed on TapeStation (Agilent). Gene expression libraries were sequenced on a Novaseq 6000 System (Illumina) and cell surface protein libraries were sequenced on the NextSeq 550 System (Illumina).

### Computational analysis

Data was aligned using cellRanger version 6.1.1 (10X Genomics) to the GRCh38 reference and the RSV genome (GCA_002815475.1). Ambient RNA was corrected using FastCAR (https://github.com/Nawijn-Group-Bioinformatics/FastCAR)[19]. Demultiplexing was performed using Seurat version 4.1.1[20] and SoupOrCell (https://github.com/wheaton5/souporcell)[21]. Integration was performed with FastMNN [22]. Cell type frequencies were calculated using scCODA (https://github.com/theislab/scCODA)[23]. Differential gene expression analysis was done using edgeR version 3.36.0[24] after generating a pseudobulk dataset per donor per condition for each cell type separately. Briefly, the counts were normalized by using the TMM method, after which, a quasi-likelihood binomial generalized log-linear (glmQL) model was applied and Benjamini-Hochberg correction was applied. For the interaction analysis, the following contrast was performed: (Asthma_RSV_-Asthma_Non-Infected_)-(Healthy_RSV_-healthy_Non-infected_). Gene ontology (GO) analysis was conducted with g:Profiler and further visualization was realized for the enriched pathways from the GO:BP database. Gene Set Enrichment Analysis (GSEA) was performed using fgsea (https://github.com/ctlab/fgsea)_1_, using the gene set collections from the Release 7.4 of the Molecular Signatures Database (MSigDB). Cell-cell interaction analysis was done with CellChat version 1.1.3 (https://github.com/sqjin/CellChat)[25].

### Statistical analysis

Comparisons between patient groups at a single time point were analyzed using the non-parametric Wilcoxon-rank sum test for paired observations. Comparisons between two time points within the same group were done using two-way ANOVA test, using Graph Pad Prism Version 9.3.1 (www.graphpad.com).

### Data availability

All data will be available on EGA.

## Results

### Cellular composition of primary epithelial cells differentiated in ALI cultures

To compare the cell-type specific response to RSV between pBECs obtained from patients with asthma and healthy controls, we infected pBECs from ALI cultures with RSV, and processed the cultures for scRNAseq 72hpi (Figure 1a).

**Figure 1:**
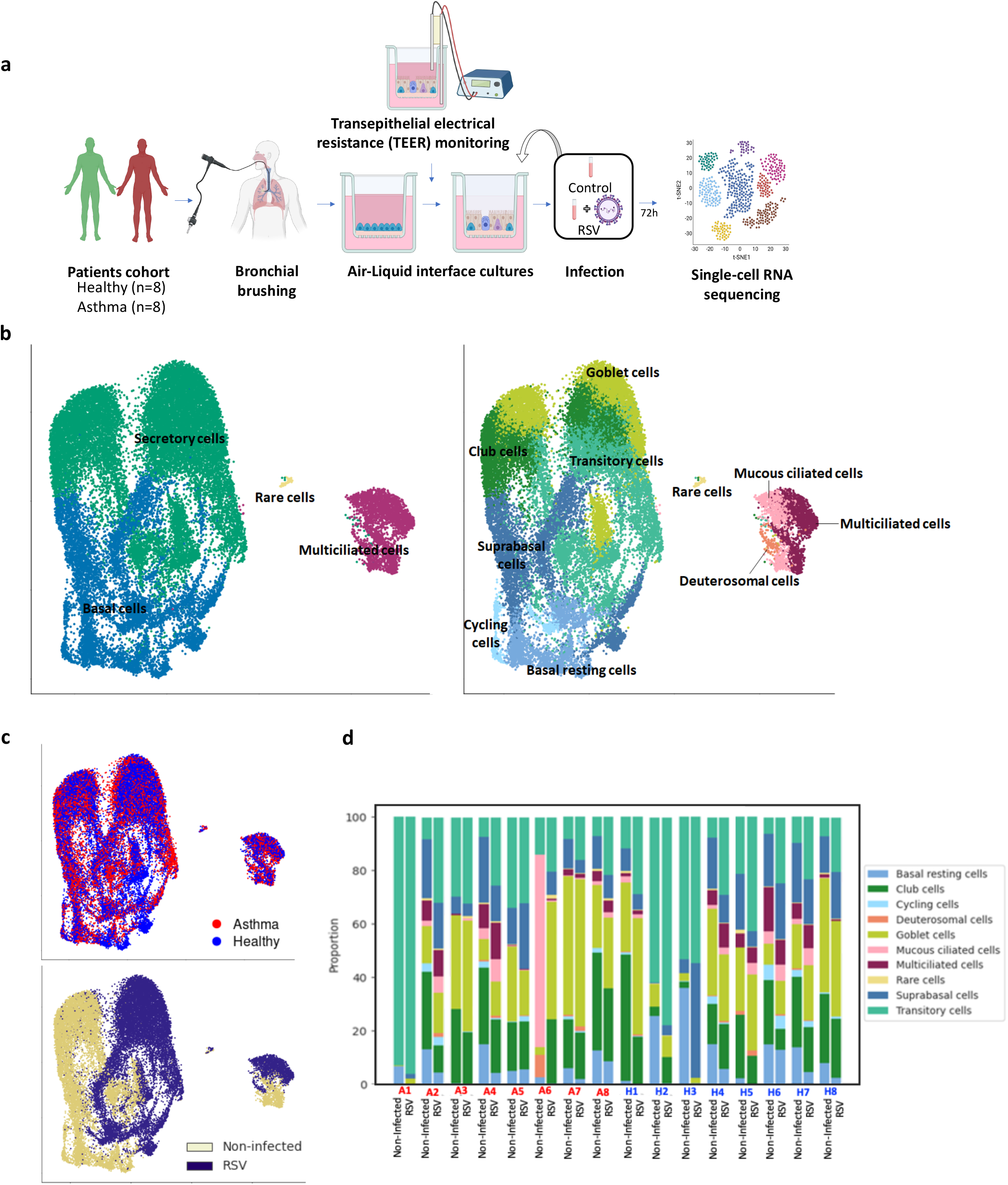
Experimental design and dataset overview. (**a)**. Overview of the study design. BECs were obtained from healthy subjects and asthma patients by bronchial brushes, and were differentiated in ALI cultures for 3 or 4 weeks before being treated with RSV. Cells were harvested and processed for scRNAseq. (**b**) UMAP representation of 30,604 single cells from all donors and conditions, clustered and annotated in 4 groups (left) and 10 sub-groups (right), (**c**) colored by disease (top) and treatment condition (bottom). (**d**) Proportions of cell types for each sample, calculated with scCODA.

In total, we generated a dataset of 30,604 cells. Unsupervised clustering identified four main epithelial cell types in the ALI cultures: basal, secretory, multiciliated and rare cells (Figure 1b). Subclustering at higher granularity allowed the identification of 10 different subsets of epithelial cells, including well-known or previously described subsets[18][26], such as suprabasal, club and goblet cells, as well as deuterosomal and mucous ciliated cells (Figure 1b). The RSV infection of the cells appeared to cause a transcriptional shift for each cell type, while the status of the donor (healthy or asthma) showed little influence on the clustering (Figure 1c). For all donors, all types of basal and secretory cells were observed, but multiciliated cells were not always present (Figure 1d).

### Cellular composition, transcriptional profile and barrier formation are similar between the ALI cultures derived from the asthma patients and the ones derived from healthy subjects

To assess whether the ALI cultures of pBECs derived from healthy subjects and asthma patients displayed any differences at baseline, we compared the transcriptomic profiles of the non-infected cells between these two groups.

For both groups, all epithelial cell subsets were identified (Figure 2a), with no differences in relative proportion (Figure 2b). Cell-type specific differential gene expression (DGE) analysis revealed no significantly differentially expressed genes (DEGs) between the two groups in any of the cell types, and no pathways were found enriched in the subsequent GSEA, indicating that the epithelial cell phenotypes were very similar. The expression level of *CX3CR1, EGFR, IGF1R, NCL*, and *ICAM1* all known to facilitate RSV entry to the cells[10,27], also showed similar expression in both groups (Supp. Figure 1). The same was observed for *DDX58* and *IFIH1*, encoding for RIG-I-like receptors, involved in the sensing of viral RNA and subsequent antiviral response[28].

**Figure 2:**
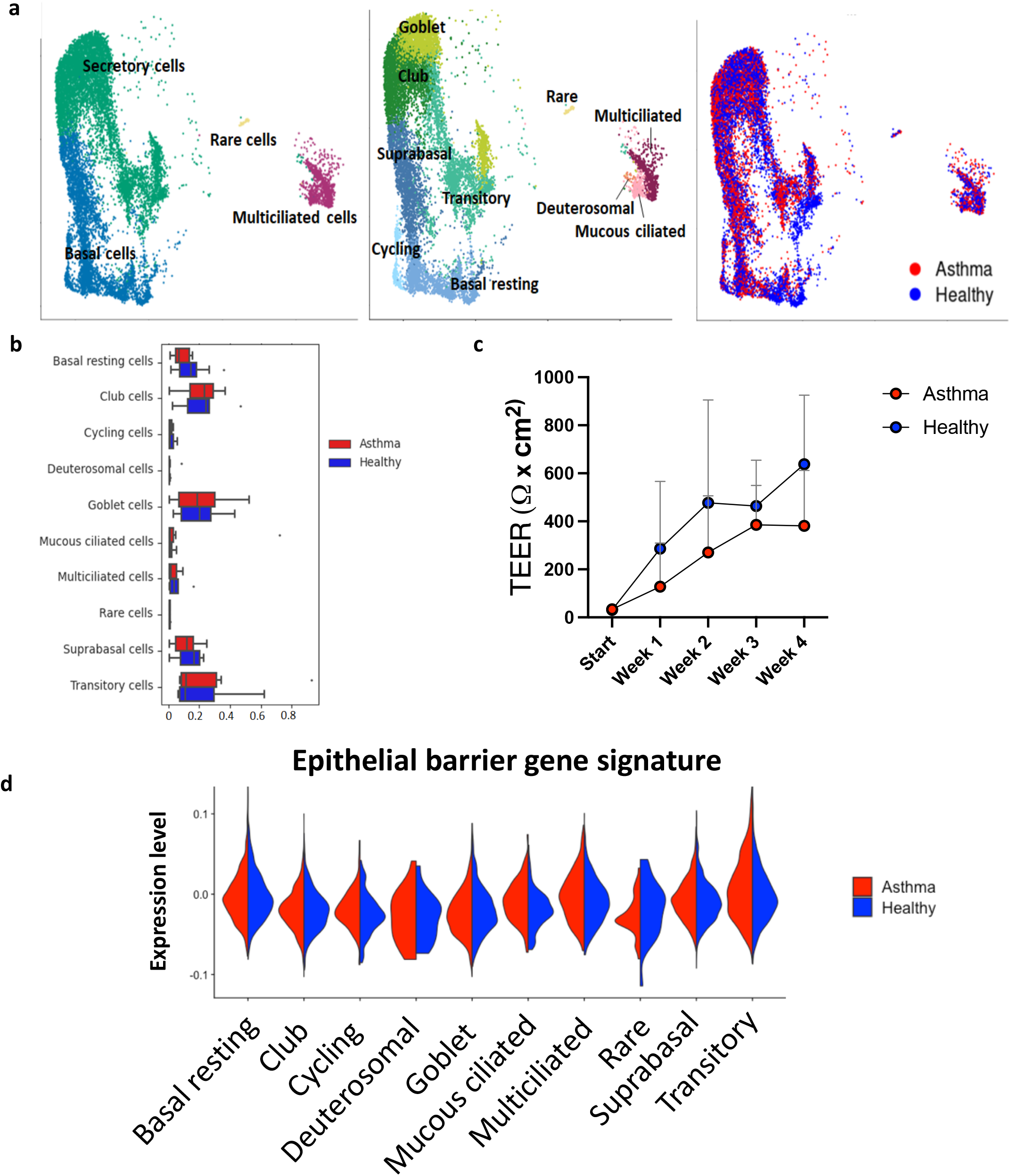
Similarities in cellular composition, barrier formation and transcriptional profiles between the ALI cultures derived from the asthma patients and the ones derived from healthy subjects. (**a**) UMAP representation of untreated 7,089 from untreated control and 6,382 asthma cells, colored by cell type (left), cell subtype (middle) and disease status (right). (**b**) Cellular frequencies of each cell subtype, colored by disease status. Significance was determined by scCODA with an FDR adjusted p-value <0.05. (**c**) TEER of the ALI cultures through time. Data are presented as mean±SD, comparisons between patient groups at a single timepoint were analyzed using the Wilcoxon-rank test. Comparisons between two time points within the same group were done using two-way ANOVA test. (**d**) Violin plots of the composite score of the genes involved in the epithelial barrier.

For both healthy- and asthma-derived ALI cultures, TEER values increased though time, suggesting the formation of an epithelial barrier. No significant difference was observed between the two groups (Figure 2c), and the expression of a signature of genes involved in the epithelial barrier[29] was also similar between asthma and healthy ALIs (Figure 2d).

### RSV infection induces a strong immune response in pBECs from healthy donors

We then investigated the response of healthy-controls derived ALI cultures of pBECs to RSV. We obtained 7,089 control treated cells and 9,199 RSV infected cells from the healthy donors (Table 2), and all 10 subtypes of epithelial cells were identified in both conditions (Figure 3a). No change in cell type proportions was observed in the samples infected with RSV compared to control treatment (Figure 3b). In addition, no changes in TEER or in expression level of a signature of epithelial barrier genes[29] were observed after RSV infection (Supp. Figure 2a,c). For each cell type, we identified the genes differentially expressed after RSV infection of the cultures by performing paired DGE analysis between untreated and infected samples (Figure 3c, Supp. Table 1). Cells transitioning from suprabasal to secretory cells had the highest number of DEGs (1,045). GO analysis of the 151 DEGs shared between basal, secretory and ciliated cells in response to RSV revealed an enrichment of biological processes, such as response to virus and cytokine production (Table 3, Supp. Table 2), indicating that the RSV infection triggered a pro-inflammatory and antiviral response from these three different types of epithelial cells.

**Table 2:** Overview of the number of cells in the scRNAseq dataset per donor per condition, for the pBECs derived from the healthy donors (top) and the asthma patients (bottom), after quality control.

| Donors (healthy) | Number of cells |  |
| --- | --- | --- |
|  | Control | RSV |
| H1 | 219 | 248 |
| H2 | 1,105 | 1,024 |
| H3 | 960 | 767 |
| H4 | 1,692 | 2,156 |
| H5 | 135 | 478 |
| H6 | 1,307 | 1,986 |
| H7 | 1,352 | 1,922 |
| H8 | 319 | 628 |
| **Total** | **7,089** | **9,199** |

| Donors (asthma) | Number of cells |  |
| --- | --- | --- |
|  | Control | RSV |
| A1 | 195 | 319 |
| A2 | 1,976 | 2,095 |
| A3 | 179 | 218 |
| A4 | 1,505 | 1,710 |
| A5 | 662 | 767 |
| A6 | 36 | 204 |
| A7 | 729 | 689 |
| A8 | 1,100 | 1,932 |
| **Total** | **6,382** | **7,934** |

**Table 3:** GO analysis of the 151 genes found differentially expressed between untreated and RSV infected samples and shared between basal, secretory and multiciliated lineages. The 25 most enriched processes are displayed.

| Biological process (GO database) | Term ID | Adjusted p-value |
| --- | --- | --- |
| Response to other organism | Go:0051707 | 9.461725873081114e-49 |
| Defense response to virus | Go:0051607 | 1.0256624683287585e-48 |
| Response to external biotic stimulus | Go:0043207 | 1.0928882461590793e-48 |
| Defense response to symbiont | Go:0140546 | 1.2024119388813918e-48 |
| Biological process involved in interspecies interaction between organisms | Go:0044419 | 4.990278548577266e-48 |
| Response to biotic stimulus | Go:0009607 | 6.60851840516966e-48 |
| Response to virus | Go:0009615 | 1.630041211843226e-47 |
| Defense response to other organism | Go:0098542 | 2.234003461280931e-46 |
| Innate immune response | Go:0045087 | 1.0338142206864037e-43 |
| Defense response | Go:0006952 | 3.994466335780032e-43 |
| Immune response | Go:0006955 | 6.899998050993308e-37 |
| Response to external stimulus | Go:0009605 | 7.385110179795586e-37 |
| Immune system process | Go:0002376 | 1.9919579659955983e-30 |
| Regulation of viral process | Go:0050792 | 7.273287357780607e-30 |
| Negative regulation of viral process | Go:0048525 | 1.083345310148469e-29 |
| Regulation of viral life cycle | Go:1903900 | 2.1337332693546325e-28 |
| Response to stress | Go:0006950 | 6.4933736039781e-27 |
| Regulation of response to biotic stimulus | Go:0002831 | 1.0780413145692529e-26 |
| Viral process | Go:0016032 | 2.2219406695936574e-24 |
| Negative regulation of viral genome replication | Go:0045071 | 1.305438220320424e-23 |
| Regulation of viral genome replication | Go:0045069 | 2.6988811960322766e-23 |
| Regulation of defense response | Go:0031347 | 3.059433362307539e-23 |
| Response to cytokine | Go:0034097 | 1.301887546010741e-22 |
| Response to type I interferon | GO:0034340 | 7.281206747097304e-22 |
| Viral life cycle | Go:0019058 | 1.3834154909898275e-21 |

**Figure 3:**
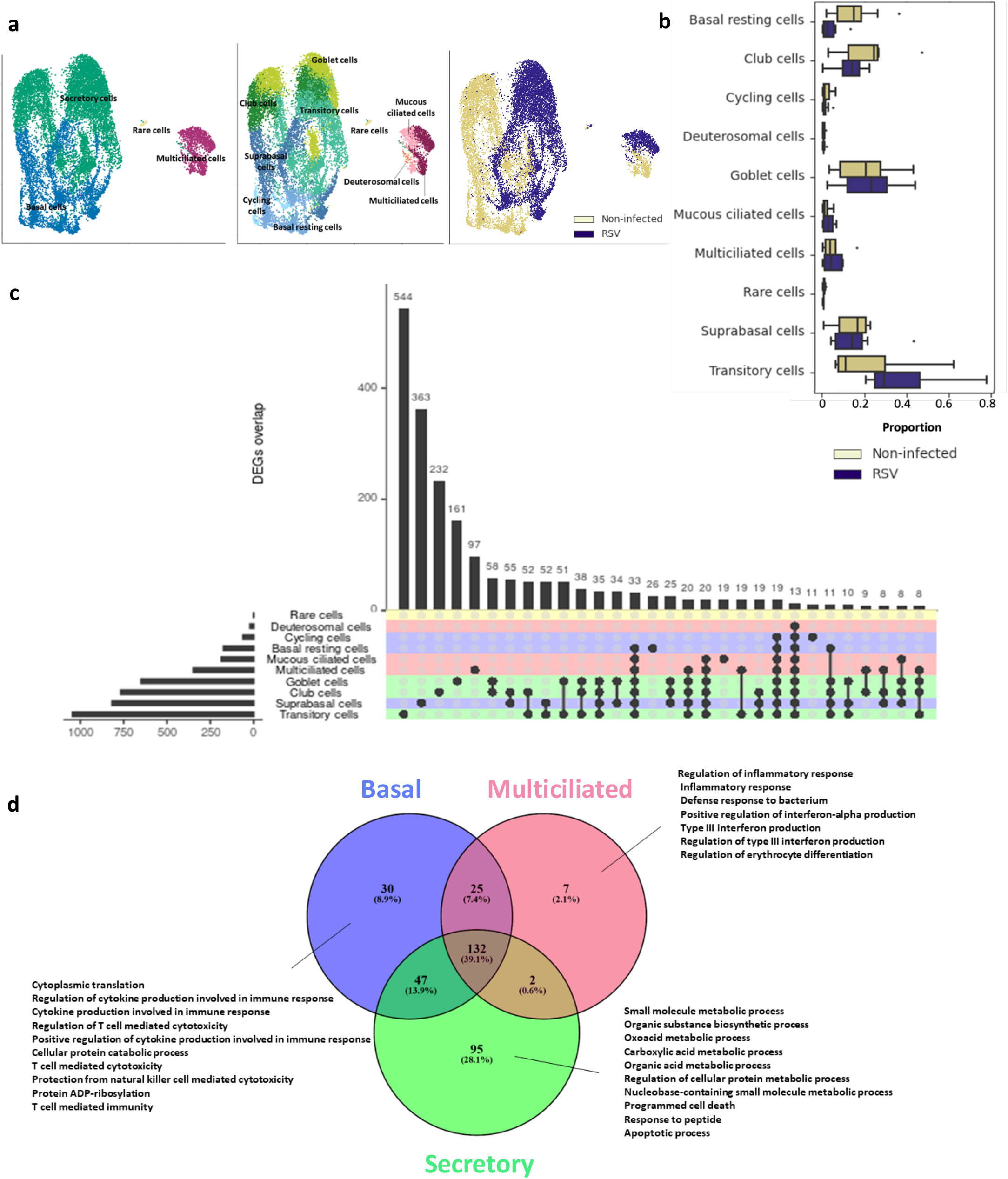
RSV infection induces transcriptomic changes in ALI cultured pBECs derived from healthy subjects. (**a**) UMAP representation of untreated 7,089 from untreated and 9,199 RSV infected cells, colored by cell subtype (left) and treatment (right). (**b**) Cellular frequencies of each cell subtype, colored by treatment. Significance was determined by scCODA with an FDR adjusted p-value <0.05. (**c**) UpSet plot depicting the unique and shared sets of DEGs with RSV infection among cell subtypes. A FDR of less than 0.05 was considered as statistically significant. (**d**) Venn diagram of the biological processes found to be significantly enriched by GO analysis of the genes differentially expressed in RSV compared to control, in basal, secretory and multiciliated cells. The 10 most enriched processes are indicated.

GO analysis of all the DEGs found for the basal, secretory and ciliated cells separately revealed an enrichment of biological processes involved in type III interferon response in response to RSV only in multiciliated cells (Figure 3d, Supp. Table 2). Biological processes related to T cell mediated immunity were exclusively found enriched in the basal cells, while many metabolic processes were only observed for the secretory cells.

Cell-cell communication analysis showed that RSV infection induced an increase in number of interactions from all cell types with each other (Supp. Figure 3a). Most cell-cell interactions were predicted to be stronger in the RSV-infected condition. We found that, compared to untreated cells, many key signaling pathways were enriched for the RSV-infected cells (Supp. Figure 3b). Among those, SEMA6, TGFb, IL1, ANGPTL, TRAIL and the DESMOSOME signaling pathways were essentially not present in the untreated condition.

### In pBECs from asthma, the RSV-induced response is largely similar to that of healthy-derived pBECs

Next, we assessed the response to RSV of the ALI cultures of pBECS derived from asthma patients. For those, all 10 subtypes of pBECs were observed (Supp. Figure 4a), without any relative changes in proportion induced by RSV (Supp. Figure 4b). Similarly to the healthy-derived cultures, no significant changes in TEER, nor in in expression of the genes involved in the barrier formation were observed 72hpi (Supp. figure 2b,c).

To investigate the transcriptional response to RSV in asthma-derived pBECs, we performed paired DGE analysis between untreated and infected samples per cell-type. For all cell types, fewer DEGs were found than in the pBECs from healthy donors (Figure 4a, Supp. Figure 4c, Supp. Table 3). However, similarly to what was observed in the healthy pBECs, the highest number of DEGs in asthma was detected in the transitory cells (Supp. Figure 4c). GO analysis of all the DEGs found for the basal, secretory and ciliated cells separately revealed an enrichment of biological processes involved immune and antiviral response for all cell types (Supp. Figure 4d, Supp. Table 4). No biological processes were found to be exclusively enriched for the multiciliated cells in asthma. However – and similarly to what was observed in healthy - many metabolic processes were only found enriched in the secretory cells, and the processes related to lymphocyte mediated immunity were exclusively being enriched in the basal cells.

**Figure 4:**
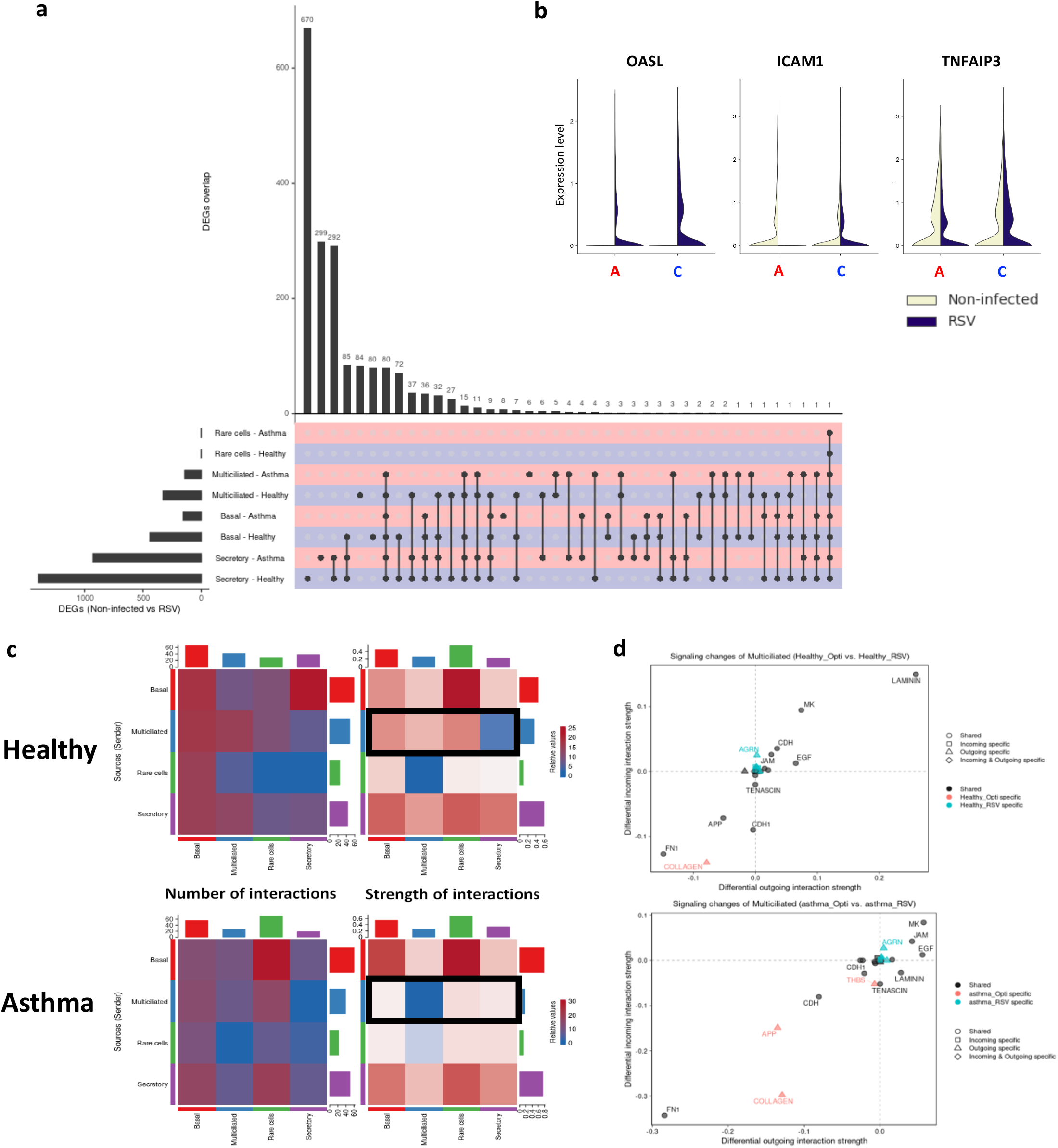
RSV infection induces a different transcriptomic response and a shift in cellular communication in pBECs derived from asthma patients compared to control. (**a**) UpSet plot depicting the unique and shared sets of DEGs with RSV infection among cell types in control (blue) and in asthma (red). A FDR of less than 0.05 was considered as statistically significant. (**b**) Violin plots of the expression of *OASL, ICAM1* and *TNFAIP3* in goblet cells derived from asthma patients (**A**) and healthy controls (**C**). (**c**) Differential number and strength of cell-cell interactions when comparing untreated to RSV infected cells derived from healthy subjects (top) and from asthma patients (bottom). Red (or blue) depicts an increase (or a decrease) of these metrics in the RSV infected cells compared to untreated cells. (**d**) Signaling changes of multiciliated cells in control (top) and in asthma (bottom) when comparing untreated and RSV infected samples.

In addition, an enrichment of major inflammatory pathways after RSV infection, including IFNα response, IFNγ response and NF-kB response, was observed similarly for the asthma and the healthy samples at the gene expression level (Supp. figure 5a). At the protein level, quantification by ELISA revealed higher concentrations of IFN-λ 1/3 in the apical washes of the RSV-infected 24hpi and 48hpi samples compared to the uninfected samples. No differences in levels of IFN-β were detected (Supp. figure 5b).

### In asthma, the expression of several genes involved in the antiviral response is altered in goblet cells

By comparing the size of the changes in gene expression induced with RSV between the healthy-derived cells and the asthma-derived cells, we observed several genes, for each cell type, displaying an opposite effect in asthma than in healthy (Supplementary figure 6). To determine which genes were differentially regulated in response to RSV infection between pBECs from patients with asthma and those from healthy controls, we performed an interaction analysis per cell type. We found a significant interaction for 8 genes in total, including 6 differentially regulated in goblet cells (Table 4). Of these, *OASL, ICAM1* and *TNFAIP3* display an impaired expression in asthma samples infected with RSV compared to control (Figure 4b), with both *OASL* and *TNFAIP3* being more induced by RSV in the healthy goblet cells than in the goblet cells from asthma donors, and *ICAM1* not being induced by RSV in the goblet cells in asthma.

**Table 4:**
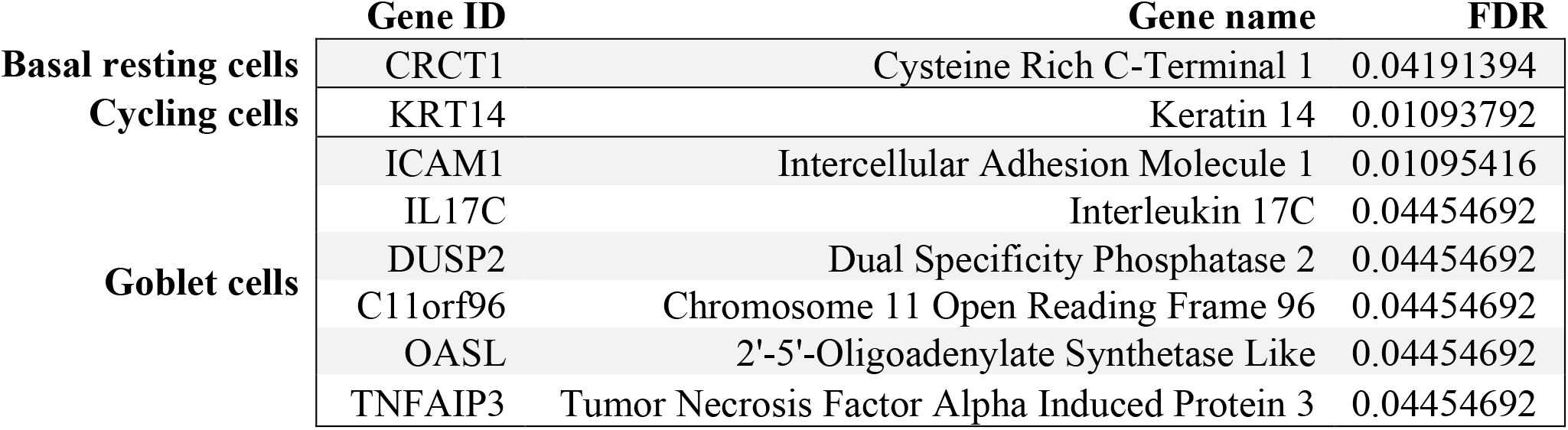
Significant genes from the interaction term (asthma_RSV_-asthma_Non-Infected_)-(healthy_RSV_- healthy_Non-Infected_). Interaction analysis was performed using edgeR, for each cell type. Statistical significance was considered for genes with a FDR<0.05.

### Change in cellular communication induced by RSV is impaired in asthma

Cell-cell interaction analysis showed similar changes of behavior with RSV in both asthma- and healthy-derived cells, with an increase in the number of interactions after RSV infection, but a decrease in the strength of those interactions (Supp. figure 7a). Signals sent from multiciliated cells to secretory cells were increased and stronger with RSV for the healthy pBECS, but were decreased in asthma (Figure 4c). For both groups, the outgoing collagen signaling from multiciliated cells was absent from the RSV infected cells (Figure 4d). Interestingly, APP (amyloid precursor protein) and THBS (thrombospondin) signaling, both found in the untreated and RSV infected healthy multiciliated cells, were only detected in the untreated condition in asthma, but not in RSV infected cells. In basal and in secretory cells, they were found in both untreated and RSV conditions (Supp. Figure 7b,c).

## Discussion

RSV plays a critical role in early life recurrent wheeze and in asthma exacerbations. The response to RSV infection differs between the healthy and the asthma bronchial epithelium. In this study, we compared the transcriptome of bronchial epithelial cells from healthy and asthma donors in ALI cultures before and after RSV infection, to better understand the molecular and cellular mechanisms causing this difference. We found a lot of similarities in the transcriptional response to RSV between pBECs of asthma patients and pBECs of healthy donors, with an overlapping antiviral response across cell types. However, the expression of genes involved in triggering the interferon response was different between goblet cells from healthy controls and patients with asthma after RSV infection, with an enrichment in type-III interferon response only found in the healthy multiciliated cells. In addition, the changes in APP and the THBS signaling observed with RSV infection in control cultures were impaired in multiciliated cells after RSV infection in cultures from patients with asthma. Overall, our study indicates that the reduced response induced by RSV in cultured airway epithelial cells in asthma are mostly observed to the most differentiated cells, suggesting that they would be a preferred target for potential drug development.

Contrarily to what others previously described, there was no reduction in TEER measurement caused by the RSV infection[30]. This could indicate that the RSV did not cause lysis of the cells in our model, and corroborates with previous studies where no obvious deterioration of ALI cultures was observed after RSV infection[31]. In similar models, it was also originally suggested that RSV would preferentially infect ciliated cells[31]. Our data suggests that it is not the case, similarly to what others reported previously[32]. However, it is important to note that for some samples of our dataset, the number of ciliated cells was limited. In addition, we didn’t observe any increase of mRNA expression encoding the viral receptors in asthma, in opposition to what was previously described[9].

In our study, the strong antiviral response after RSV infection presents similarities with the response to SARS-CoV-2 infection[33], with goblet cells showing a strong inflammatory signature in both cases. *ICAM1* and *TNFAIP3*, known to be involved in the antiviral response, were previously identified as some of the best drug targets for COVID-19[34]. In our study, *ICAM1* and *TNFAIP3* have an altered expression in goblet cells in asthma in response to RSV. The relevance of studying goblet cells in respiratory viral infections has already been demonstrated for rhinovirus, multiple influenza viruses and hantavirus[35]. Recently, goblet cells were also found to play a critical role in SARS-CoV-2 infection in the lung, and an increased viral replication in the COPD airway epithelium was observed, likely due to COPD associated goblet cell hyperplasia[36]. Our findings, taken together with the emerging role of goblet cells observed in recent literature, suggest that goblet cells play a critical role in RSV-induced asthma exacerbations. This highlights the need for deeper investigations of goblet cells, especially, in the context of airway diseases, frequently associated with goblet cells metaplasia and hyperplasia and for which antiviral response is impaired.

Last, no enrichment of type-III interferon response from the multiciliated cells was observed in asthma. Previous studies have showed that type-III interferon response was strongly induced by respiratory viruses in nasal epithelial cells [37] highlighting that an impairment of this response might contribute to the defective antiviral response in asthma. However in the context of asthma, major differences were observed between *in vivo* studies and *in vitro* studies performed in isolated bronchial epithelial cells: with most *in vivo* studies showing an antiviral IFNλ induction in asthma, while, similarly to our own results, majority of the *in vitro* studies report a defect of IFNλ[38]. The current increase of studies on the type-III interferon response to respiratory viruses should unveil in the coming years the potential benefic effect of altering the IFNλ response and impact on patients.

Overall, gaining insights at a cell type specific level about the link between the epithelium and the activities of the immune cells seems to remain necessary for future development towards RSV-mediated asthma exacerbation.

## Acknowledgements

The authors thank Uilke Brouwer, Sharon Brouwer, Marnix Jonker, Marissa Wisman and Jelmer Vlasma, from the Department of Pathology and Medical Biology of the University Medical Center Groningen, for their support in the experimental procedure. They also acknowledge Marijn Berg (University Medical Center Groningen), Alen Faiz (University of Technology Sydney), and Hana Aliee (Helmholtz Zentrum München) for their advice for conducting the bioinformatics analysis.

## Statement of Ethics Approval

The presented study did not involve human participants, nor animal subjects. All data is derived from human material, but does not allow participants identification. This material was collected from a study previously described and which obtained ethics approval and involved informed patients consent (see reference [18] for details). The ethical approval by the Institutional Review Board (IRB; METc) from this previous study (ABR n°NL53173.042.015 “A better understanding of molecular mechanisms leading to asthma and its remission (ARMS)”, and ABR n° NL69765.042.19 (“Towards Targeting the Origin of Asthma (ORIENT)”) included the use of the patients-derived cells for this type of investigations.

## Competing interests

G.H.K, M.C.N. and M.v.d.B. received project funding from GlaxoSmithKline.

G.H.K and M.v.d.B. received funding from Astra Zeneca.

M.v.d.B. received funding from Novartis, Genentech and Roche.

## Funding

Unrestricted research grant received from the Netherlands Lung Foundation (grants n°4.1.18.226 and n°5.1.14.020), as well as unrestricted research grant received from European Union’s H2020 Research and Innovation Program under grant agreement no. 874656 were used for this study. The study sponsors had no specific role in the study design; in the collection, analysis and interpretation of the data; in the writing of the report; nor in the decision to submit the paper for publication.

