## Supplemental Figures for "Airway epithelial response to RSV is impaired in multiciliated and goblet cells in asthma"

### Supplementary figures

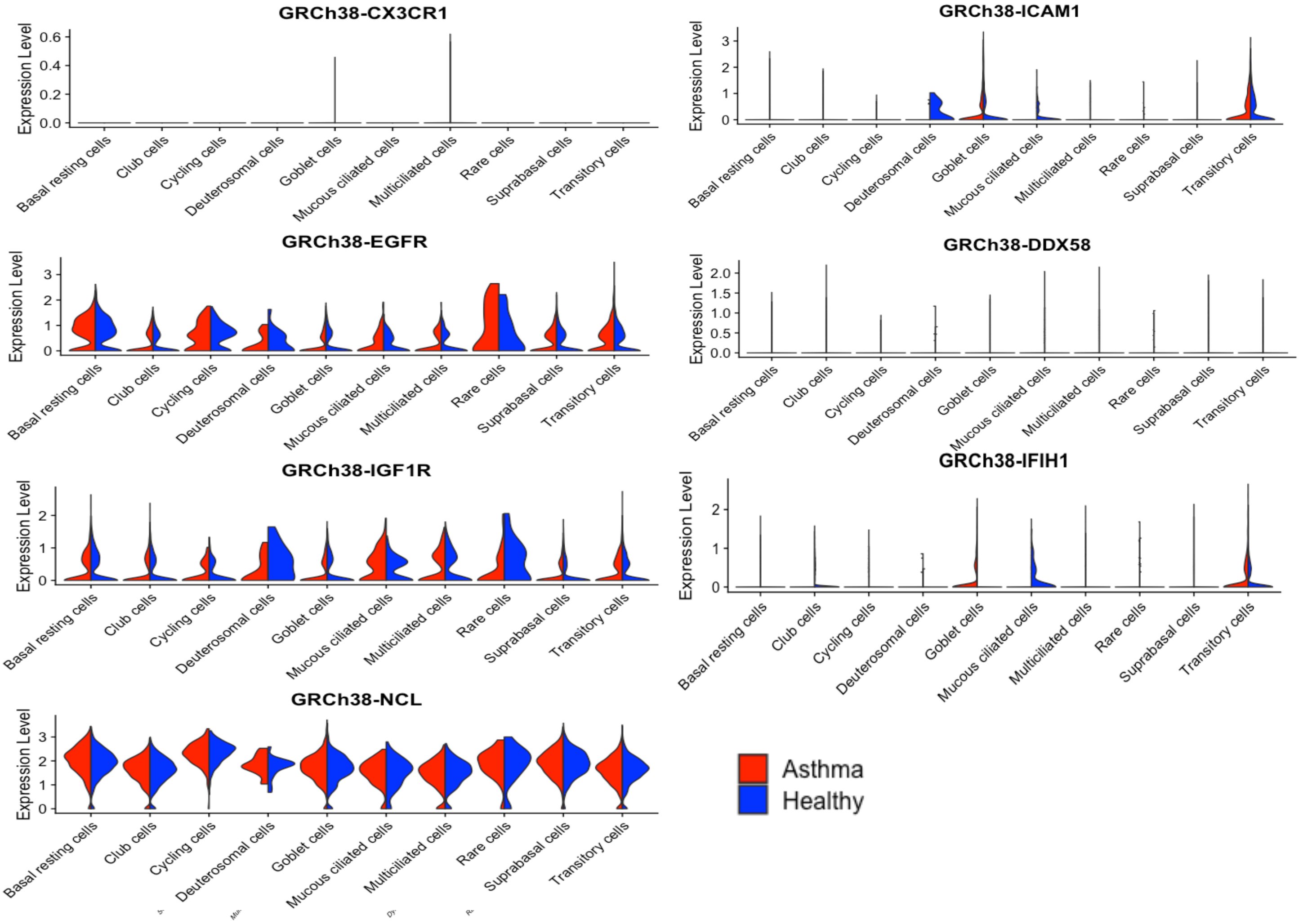


**Supplementary figure 1: ALI cultures of pBECs derived from healthy donors and asthma patients show similar expression of the RSV receptors.** Violin plots of the expression of RSV receptors for the non-infected samples. Differential gene expression between the two groups was performed in each cell type. A FDR of less than 0.05 was considered statistically significant.

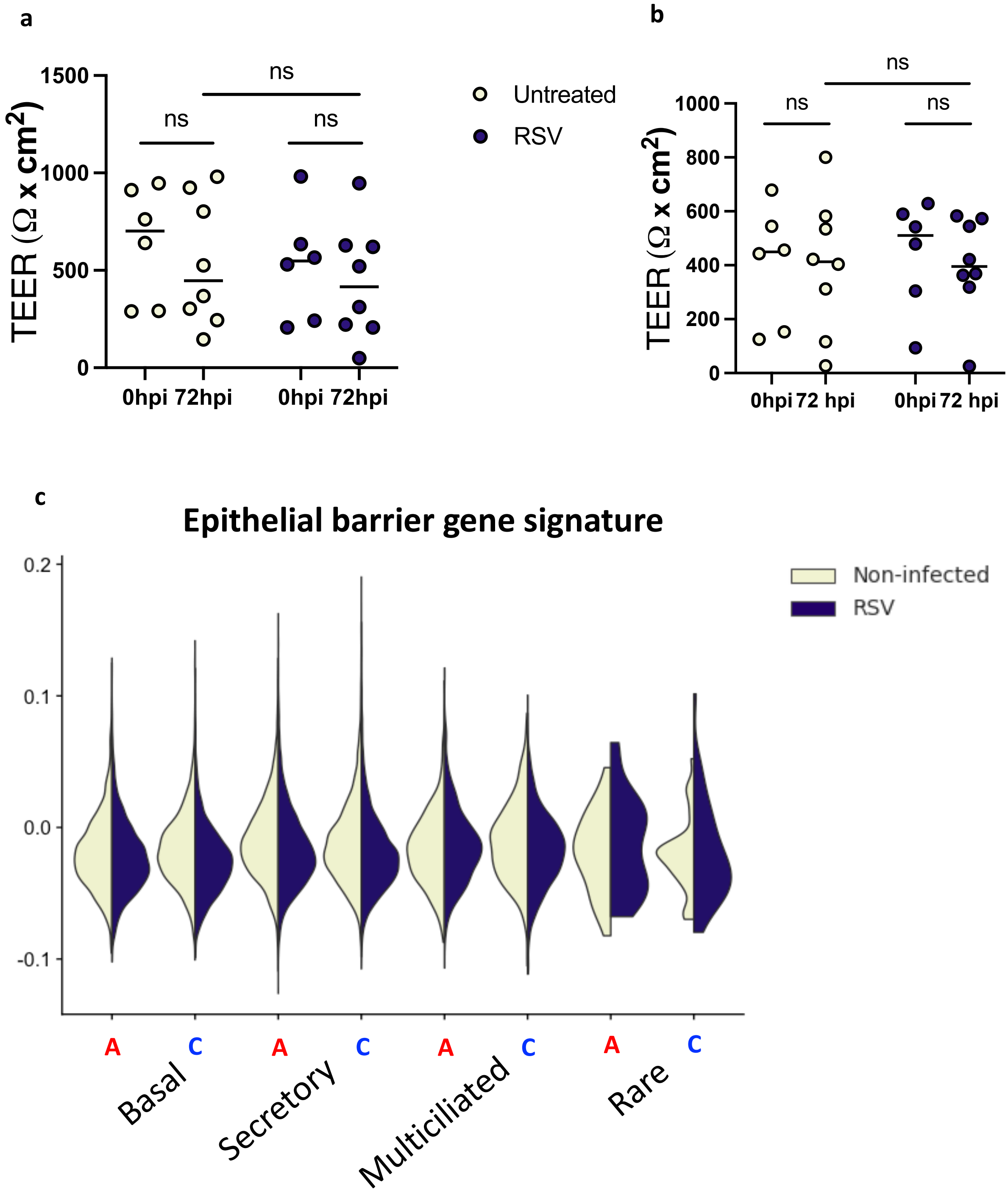


**Supplementary figure 2: RSV infection does not alter the epithelial barrier.** Comparison at 72hpi of the TEER measurements of the untreated and the RSV infected ALI cultures of pBECS derived from (**a**) control and (**b**) asthma donors. (**c**) Violin plots of the composite score of the genes involved in the epithelial barrier formation in the pBECs derived from asthma patients (**A**) and healthy controls (**C**).

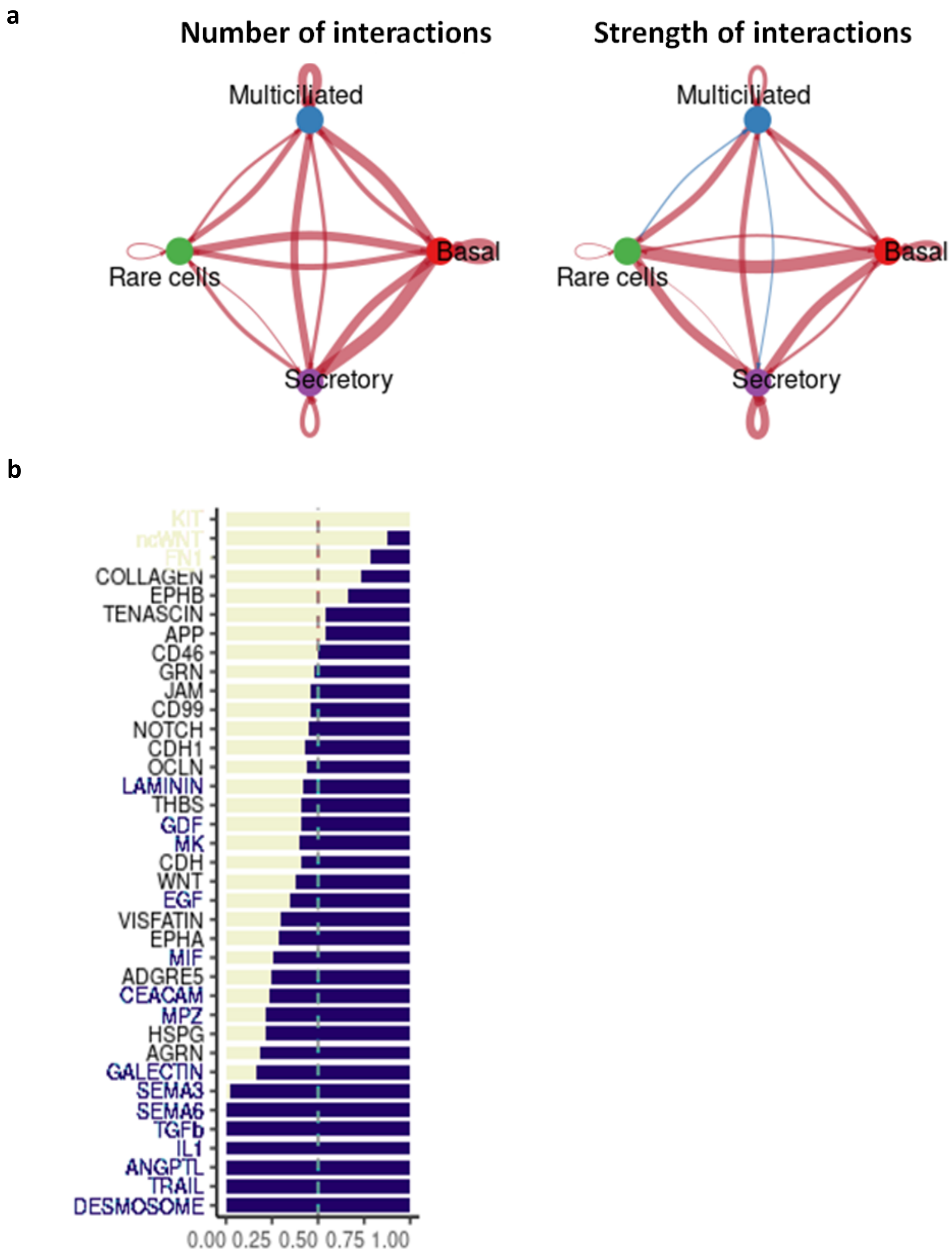


**Supplementary figure 3: RSV induces a shift in cell-cell communication in the healthy-derived pBECs.** (**a**) Differential number (left) and strength (right) of cell-cell interactions, when comparing untreated to RSV infected cells. Nodes indicate cell type and edges thickness represents the relative interaction count or interaction strength. Edges colored in red (or blue) depicts an increase (or a decrease) of these metrics in the RSV infected cells compared to untreated cells. (**b**) Significant signaling pathways (right) ranked based on differences in the overall information flow within the inferred networks, when comparing untreated to RSV infected cells. Pathways in beige (or blue) indicate a significant enrichment in the non-infected (or RSV) condition, determined by CellChat.

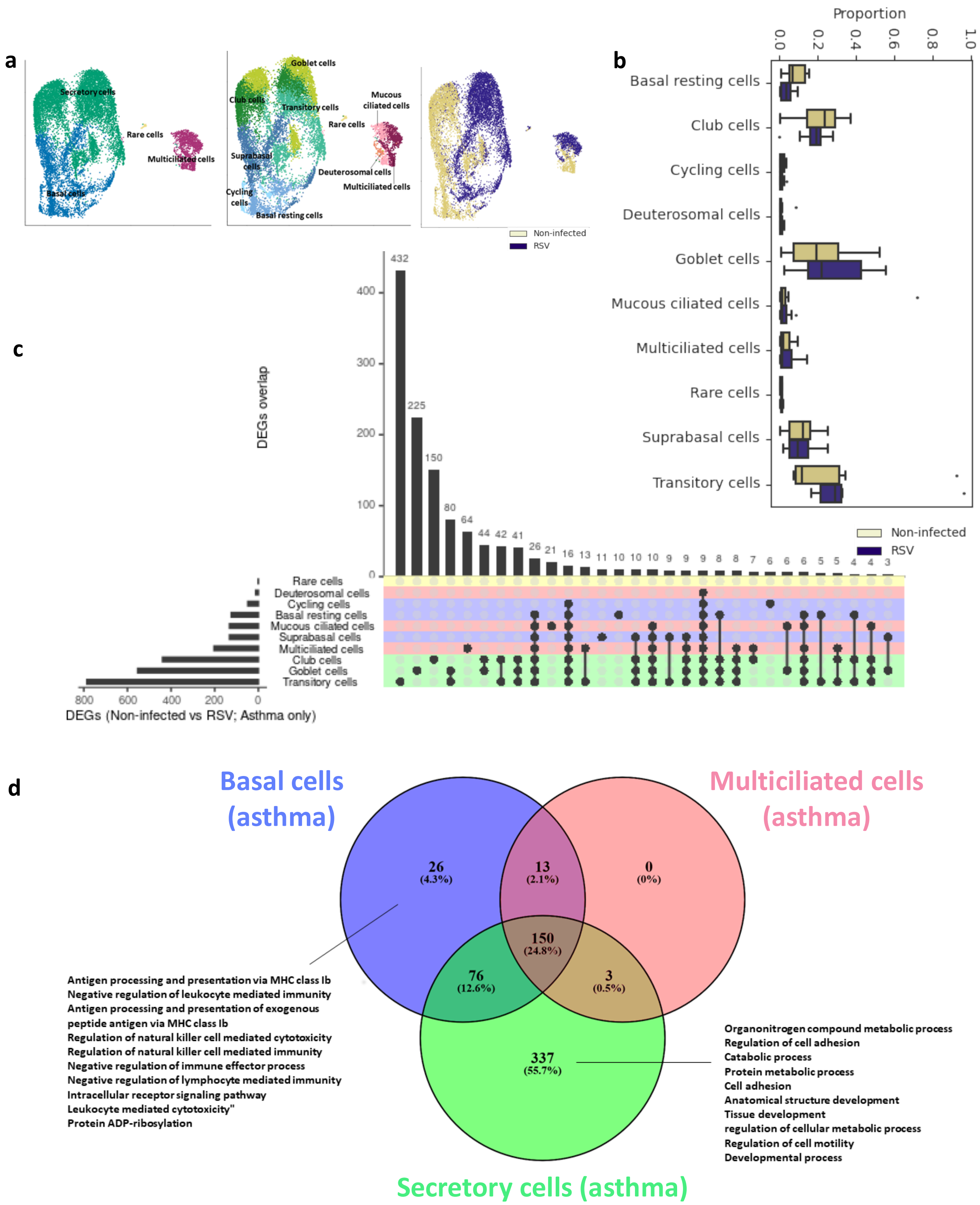


**Supplementary figure 4: RSV response in asthma** (**a**) UMAP representation of untreated 6,382 from untreated and 7,934 RSV-infected cells, colored by cell type (left), subtype (middle) and treatment (right). (**b**) Cellular frequencies of each cell subtype, colored by treatment. Significance was determined by scCODA with an FDR adjusted p-value <0.05. (**c**) UpSet plot depicting the unique and shared sets of DEGs with RSV infection among cell types. A FDR of less than 0.05 was considered as statistically significant. (**d**) Venn diagram of the biological processes found to be significantly enriched by GO analysis of the genes differentially expressed in RSV compared to control, in basal, secretory and multiciliated cells. The 10 most enriched processes are indicated.

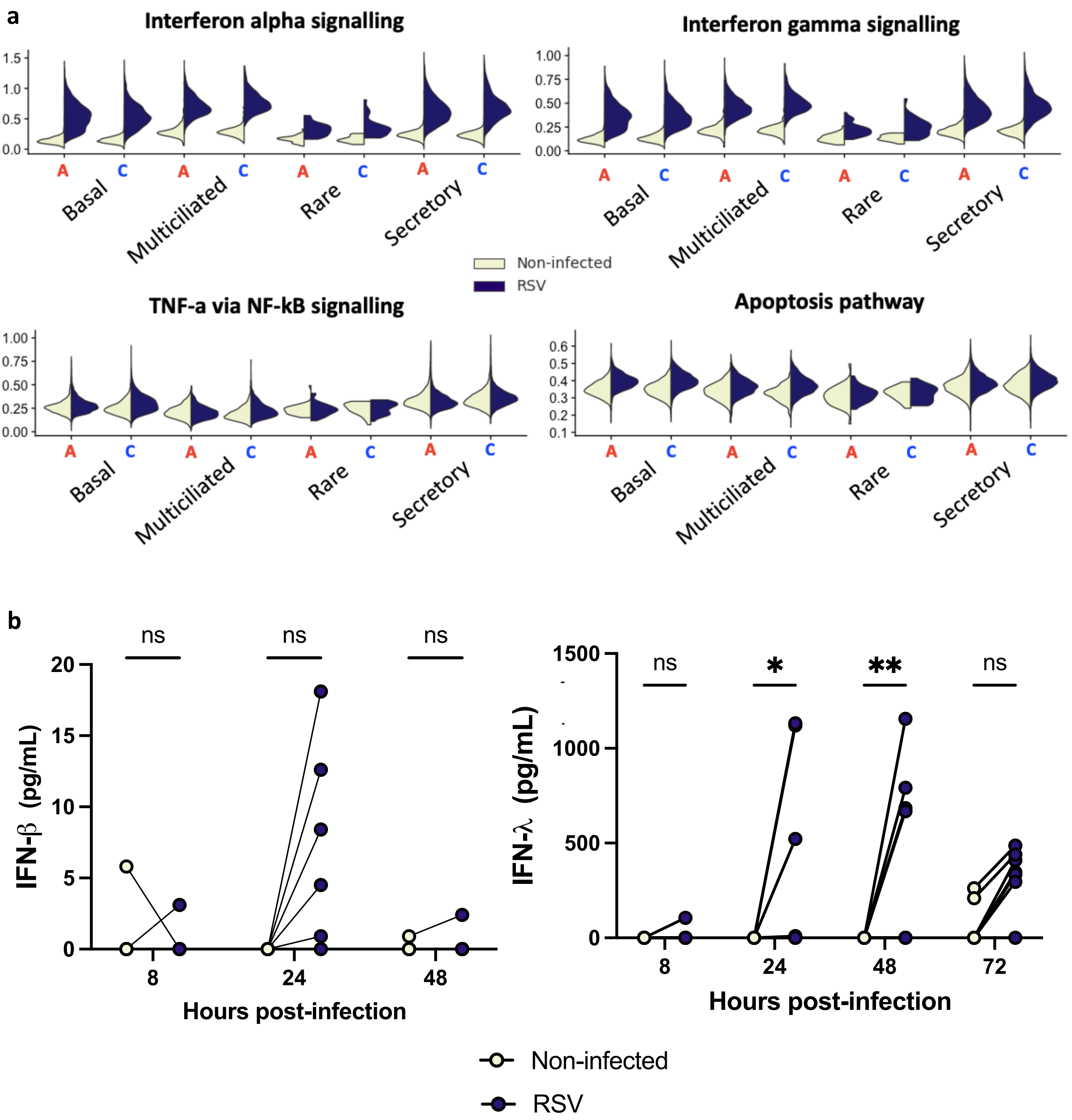


**Supplementary figure 5: RSV infection induces an antiviral response in ALI cultures of pBECs derived from both control and asthma donors.** (**a**) Violin plots of the composite score of the main antiviral response pathways in the pBECs derived from asthma patients (**A**) and healthy controls (**C**). (**b**) Concentration of IFN-ß and IFN-λ 1/3 measured by ELISA in the apical washes at different time points after infection for all samples. *: $p<0.05$, **: $p<0.01$ (between the indicated values, as analysed by the two way ANOVA test).

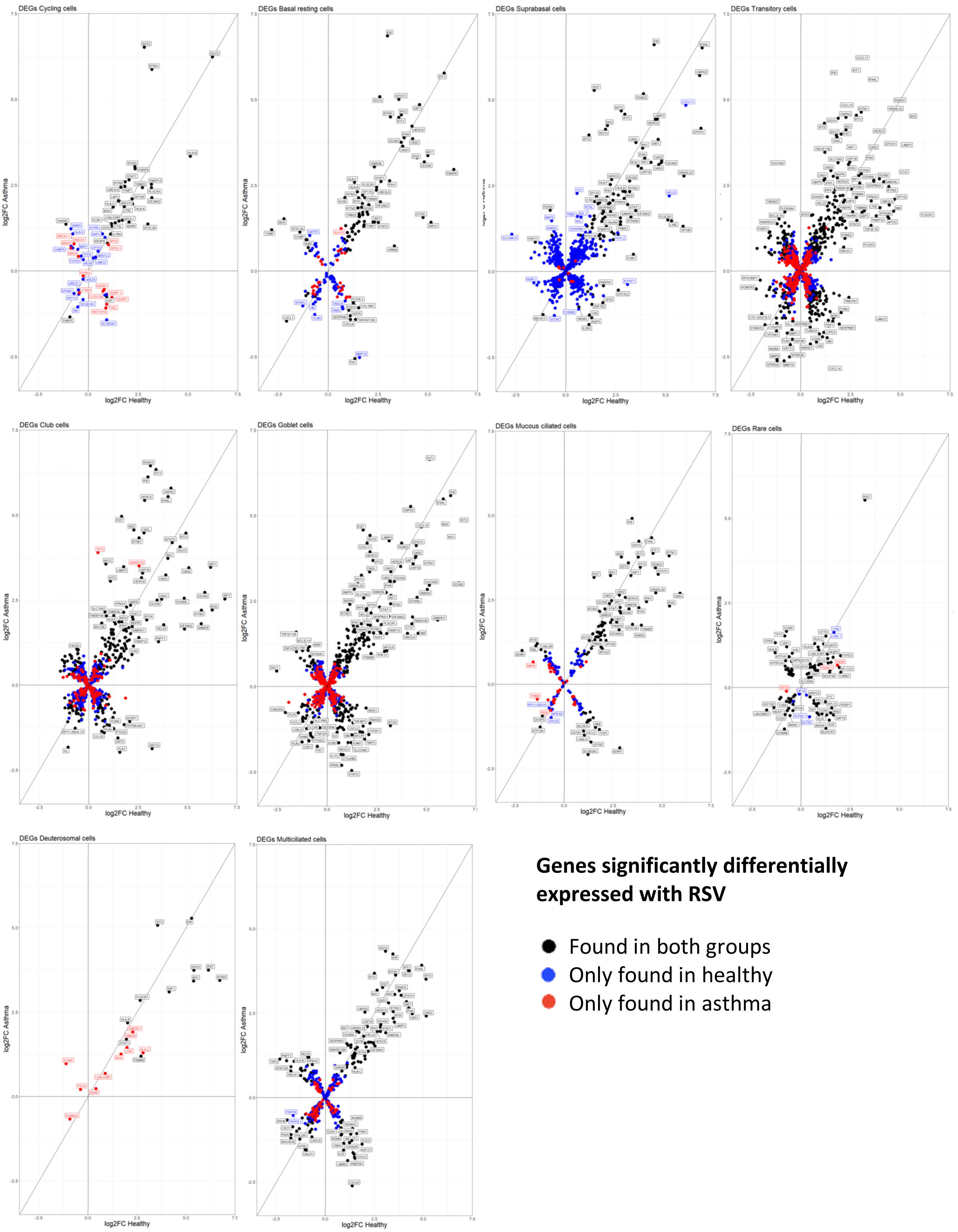


**Supplementary figure 6: Effect of RSV on gene expression of pBECs derived from asthma patients compared to pBECs derived from healthy donors.** The logFoldChange (log2FC) for the change in gene expression induced by RSV in healthy vs in asthma, for each of the 10 cell types is represented. Only genes found to be significanty different in RSV (FDR<0.05) in either the healthy- or the asthma-derived pBECs are displayed. DEGs found in both conditions are colored in **black**, DEGs only found in healthy (resp. asthma) are colored in **blue** (resp. **red**).

**a**

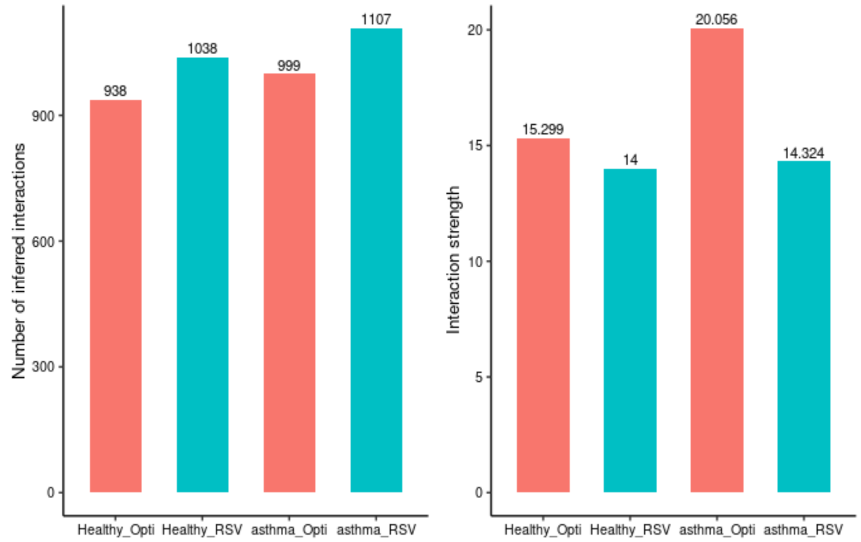
Number of inferred interactions
938
1038
999
1107
Healthy_Opti
Healthy_RSV
asthma_Opti
asthma_RSV
Interaction strength
15.299
14
20.056
14.324

**b**

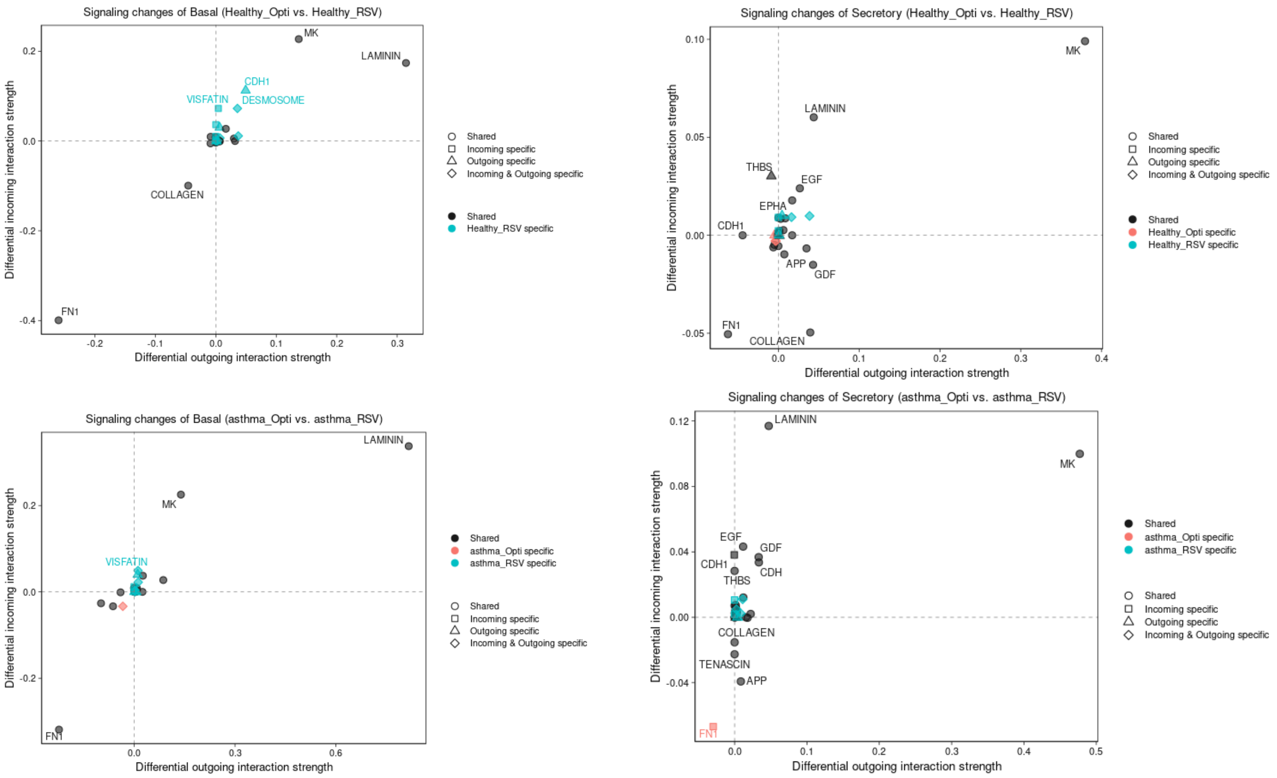
Signaling changes of Basal (Healthy_Opti vs. Healthy_RSV)
Signaling changes of Secretory (Healthy_Opti vs. Healthy_RSV)
Signaling changes of Basal (asthma_Opti vs. asthma_RSV)
Signaling changes of Secretory (asthma_Opti vs. asthma_RSV)
Differential incoming interaction strength
Differential outgoing interaction strength
Shared
Incoming specific
Outgoing specific
Incoming & Outgoing specific
Healthy_Opti specific
Healthy_RSV specific
asthma_Opti specific
asthma_RSV specific
MK
LAMININ
CDH1
VISFATIN
DESMOSOME
COLLAGEN
FN1
THBS
EGF
EPHA
CDH1
APP
GDF
CDH
TENASCIN

c

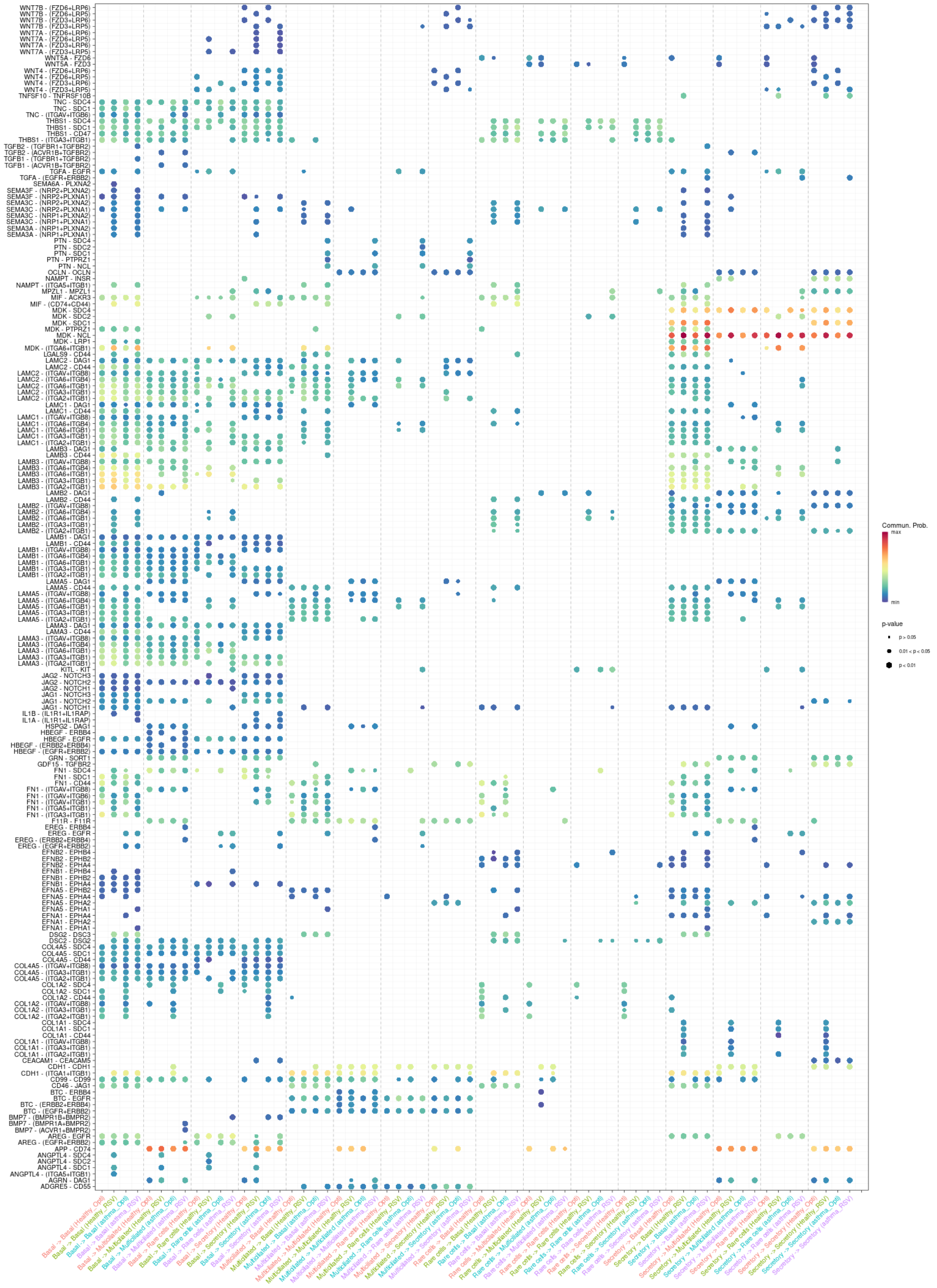


**Supplementary figure 7: Cell-cell communication changes induced by RSV differ between asthma and control.** (**a**) Number (left) and strength of inferred cellular interaction in untreated and RSV infected cells in asthma- and healthy-derived pBECs, determined with CellChat. (**b**) Signaling changes of basal (left) and secretory (right) in control (top) and in asthma (bottom) when comparing untreated and RSV infected samples. (**c**) Communication probabilities mediated by ligand-receptor pairs from some cell groups to other cell group.
